# Phylo-Node: a molecular phylogenetic toolkit using Node.js

**DOI:** 10.1101/075101

**Authors:** Damien M. O’Halloran

## Abstract

**Background:** Node.js is an open-source and cross-platform environment that provides a JavaScript codebase for back-end server-side applications. JavaScript has been used to develop very fast and user-friendly front-end tools for bioinformatic and phylogenetic analyses. However, no such toolkits are available using Node.js to conduct comprehensive molecular phylogenetic analysis.

**Results:** To address this problem, I have developed, *phylo-node*, which was developed using Node.js and provides a stable and scalable toolkit that allows the user to perform diverse molecular and phylogenetic tasks. phylo-node can execute the analysis and process the resulting outputs from a suite of software options that provides tools for read processing and genome alignment, sequence retrieval, multiple sequence alignment, primer design, evolutionary modeling, and phylogeny reconstruction. Furthermore, phylo-node enables the user to deploy server dependent applications, and also provides simple integration and interoperation with other Node modules and languages using Node inheritance patterns, and a customized piping module to support the production of diverse pipelines.

**Conclusions:** phylo-node is open-source and freely available to all users without sign-up or login requirements. All source code and user guidelines are openly available at the GitHub repository: https://github.com/dohalloran/phylo-node

## BACKGROUND

The cost of whole genome sequencing has plummeted over the last decade and as a consequence, the demand for genome sequencing technology has risen significantly [1]. This demand has meant that producing large complex datasets of DNA and RNA sequence information is common in small research labs, and in terms of human health this boom in sequence information and precipitous drop in sequencing costs has had a direct impact in the area of personalized medicine [2–5]. However, once the sequence information becomes available, perhaps the greater challenge is then processing, analyzing, and interpreting the data. To keep pace with this challenge, the development of new, fast, and scalable software solutions is required to visualize and interpret this information.

JavaScript is a lightweight programming language that uses a web browser as its host environment. JavaScript is cross-platform and supported by all modern browsers. Because JavaScript is client-side, it is very fast as it doesn't have to communicate with a server and wait for a response in order to run some code. Web browsers are ubiquitous and require no dependencies to deploy and operate, and so JavaScript represents an obvious solution for visualizing sequence information. Front-end developments using JavaScript have proven to be extremely efficient in providing fast, easy-to-use, and embeddable solutions for data analysis [6–14]. A very active community of developers at BioJS (http://www.biojs.io/) provides diverse components for parsing sequence data types, data visualization, and bioinformatics analysis in JavaScript [6,7,15–19].

Node.js provides server-side back-end JavaScript. Node.js is written in C, C++, and JavaScript and uses the Google Chrome V8 engine to offer a very fast cross-platform environment for developing server side Web applications. Node is a single-threaded environment, which means that only one line of code will be executed at any given time; however, Node employs non-blocking techniques for I/O tasks to provide an asynchronous ability, by using *callback* functions to permit the parallel running of code. Node holds much potential for the bioinformatic analysis of molecular data. A community of Node developers provides modules for bioinformatic sequence workflows (http://www.bionode.io/) which in time will likely parallel the BioJS community for the number of modules versus components. However, as of now there are no robust tools for phylogenetic analysis pipelines currently available using the Node.js codebase. To fill this void I have developed, *phylo-node*, which provides a Node.js toolkit that provides sequence retrieval, primer design, alignment, phylogeny reconstruction and as well as much more, all from a single toolkit. *phylo-node* is fast, easy to use, and offers simple customization and portability options through various inheritance patterns. The Node package manager, *npm*(https://www.npmjs.com/), provides a very easy and efficient way to manage dependencies for any Node application. phylo-node is available at GitHub (https://github.com/dohalloran/phylo-node), npm (https://www.npmjs.com/package/phylo-node), and also BioJS (https://www.npmjs.com/package/phylo-node).

## IMPLEMENTATION

Phylo-Node was developed using the Node.js codebase. The phylo-node core contains a base wrapper object that is used to prepare the arguments and directory prior to program execution. The base wrapper module is contained within the *Wrapper_core* directory (Figure 1). An individual software tool can be easily accessed and executed by importing the module for that tool so as to get access to the method properties on that object. These method properties are available to the user by using the *module.exports* reference object. Inside a driver script file, the user can import the main module object properties and variables by using the *require* keyword which is used to import a module in Node.js. The *require* keyword is actually a global variable, and a script has access to its context because it is wrapped prior to execution inside the *runInThisContext* function (for more details, refer to the Node.js source code: https://www.npmjs.com/package/phylo-node). Once imported, the return value is assigned to a variable which is used to access the various method properties on that object. For example: a method property on the *phyml* object is *phyml.getphyml()*, which invokes the *getphyml* method on the *phyml* object to download and decompress the PhyML executable. For a complete list of all methods, refer to the *README*. *md* file at the GitHub repository (https://github.com/dohalloran/phylo-node/blob/master/README.md). In order to correctly wrap and run each executable, new shells must be spawned so as to execute specific command formats for each executable. This was achieved by using *child.process.exec*, which will launch an external shell and execute the command inside that shell while buffering any output by the process. Binary files and executables were downloaded and executed in this manner and the appropriate file and syntax selected by determining the user's operating system. phylo-node was validated on Microsoft Windows 7 Enterprise ver.6.1, MacOSX El Capitan ver. 10.11.5, and Linux Ubuntu 64-bit ver. 14.04 LTS.

**Figure 1.**
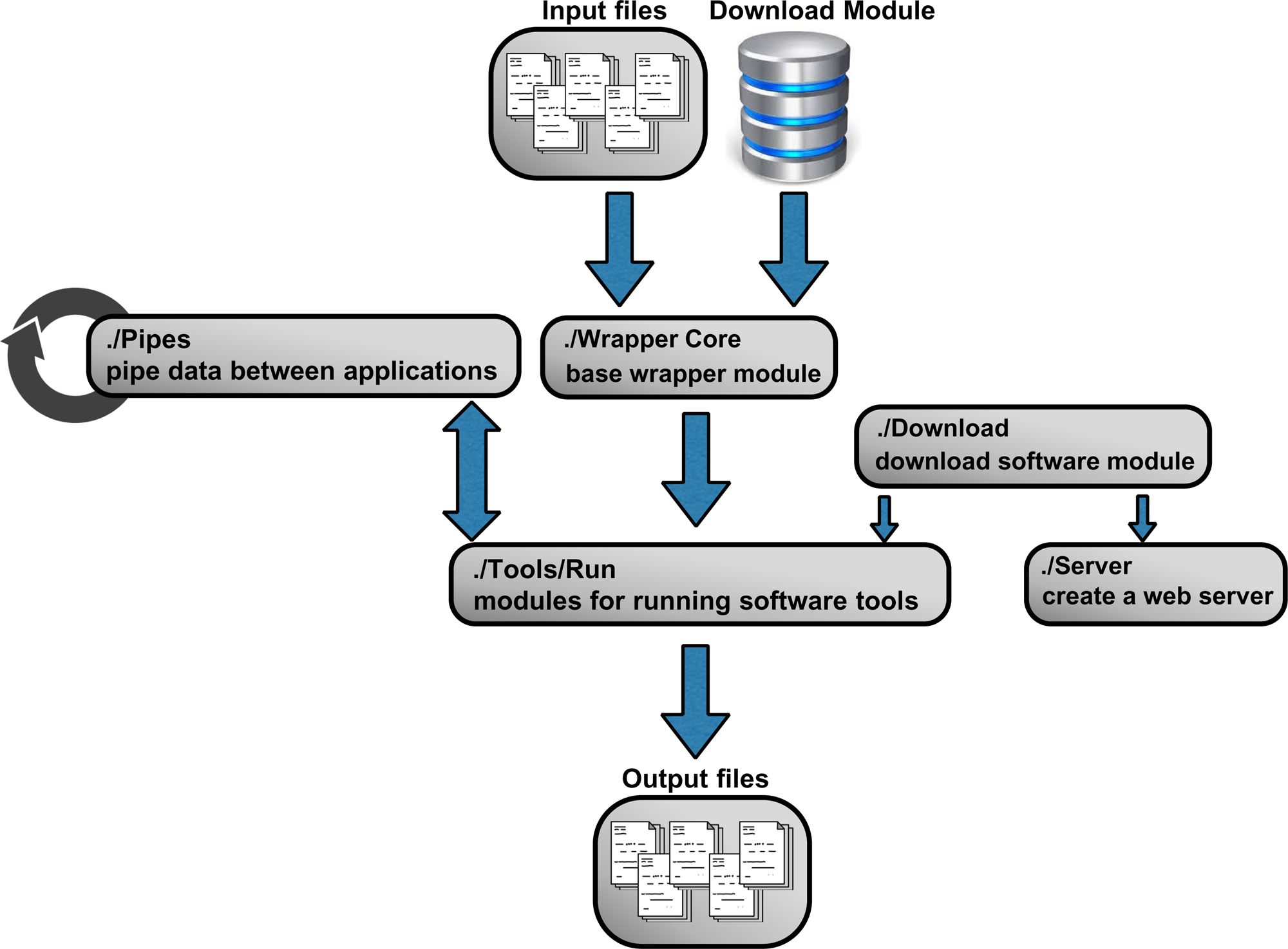
Workflow for Phylo-Node. Phylo-Node is organized into a workflow of connected modules and application scripts. In order to interface with a software tool, the base wrapper module is invoked to process command-line requests that are then passed into the software specific module. The input for the specific software can be passed into the base wrapper from a folder specified by the user or by using the sequence retrieval module which is contained within the *Sequence* directory. The *Pipes* directory contains a module for easy piping of data between applications while binaries and executables can be downloaded using the *get_executable* module from within the *Download* folder to deploy software specific modules within the *Run* directory or to provide applications to a web server from within the *Server* directory.

## RESULTS AND DISCUSSION

Phylo-Node is a toolkit to interface with key applications necessary in building a phylogenetic pipeline (Table 1). Firstly, phylo-node allows the user to remotely download sequences by building a unique URL and passing this string to the NCBI e-utilities API (http://www.ncbi.nlm.nih.gov/books/NBK25501/). Any number of genes can be supplied as command-line arguments to phylo-node by accessing the *fetch_seqs.fasta* method on the *fetch_seqs* object in order to retrieve sequence information in FASTA format. The module for remote sequence retrieval is contained within the *Sequence* directory. phylo-node also provides methods on specific objects to download various executable files using the *download* module (Figure 1). Any binary can be downloaded using the base module *get_executable* contained within the *Download* directory, however objects pertaining to specific tools such as PhyML also contain methods for downloading and unpacking binaries (see README.md file for details). phylo-node also enables the user to create a web server to deploy embeddable applications such as JBrowse [11] which provides genome visualization from within a web browser. To facilitate interoperation between various applications and components, the phylo-node package also contains a module called *phylo-node_pipes* inside the *Pipes* directory. The *phylo-node_pipes* module allows the user to easily pipe data between different applications by requiring the *child_process* module which provides the ability to spawn child processes. Through *phylo-node_pipes*, the user can chain commands together that will be executed in sequence to build consistent, and extensive pipelines. The *Pipes* directory contains sample driver scripts for using the *phylo-node_pipes* module.

**Table 1.**
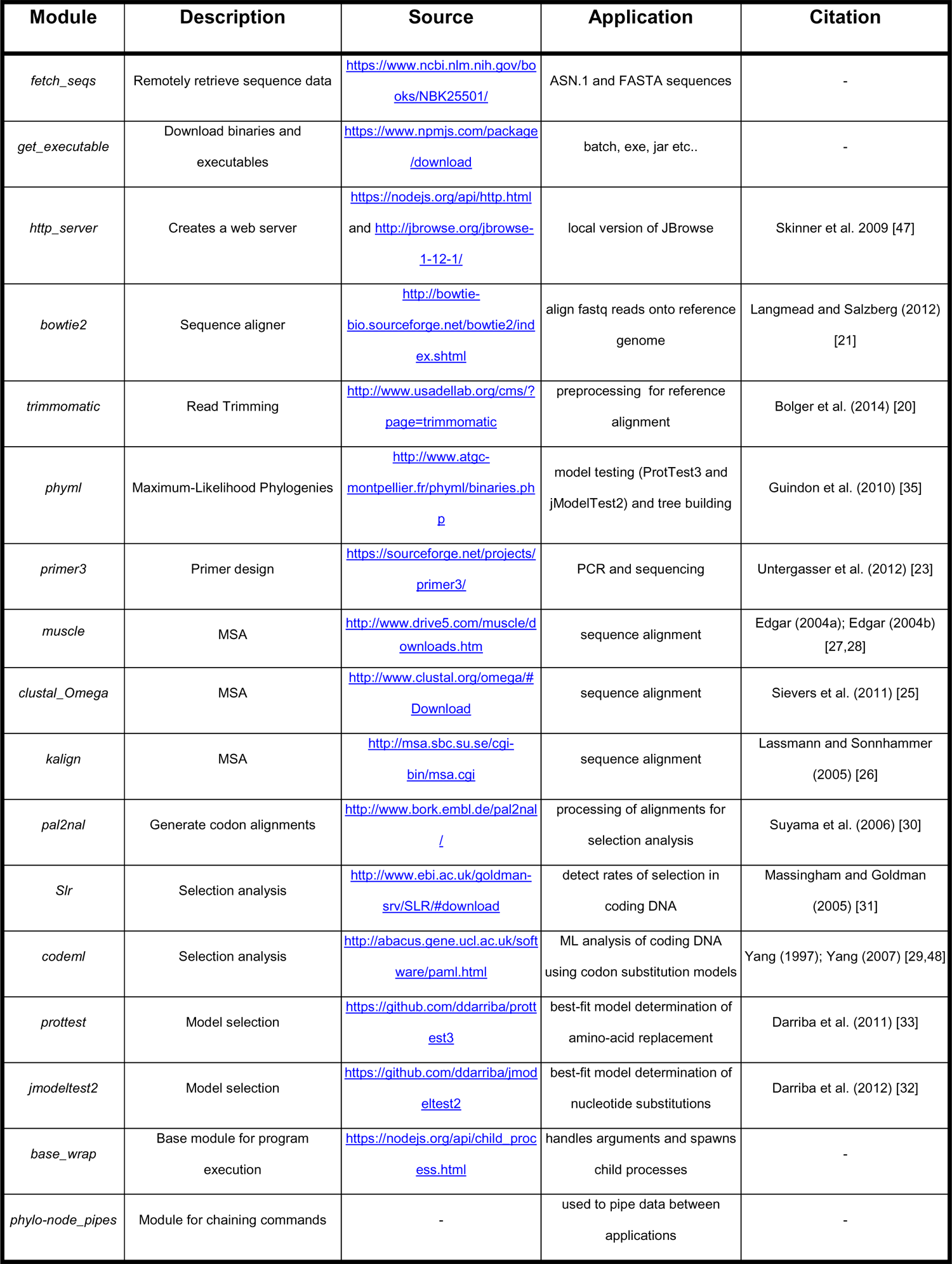
Summary of *phylo-node* applications.

Phylo-Node is highly scalable and new modules for diverse applications can easily be plugged in. The modules required to wrap and execute applications are all contained within the *Run* directory. The following tools can be implemented using phylo-node from within the *./Tool/Run* directory: Trimmomatic [20] to process reads prior to read alignment; Bowtie2 [21] for read alignment to a reference sequence;

Primer3 [22–24] to facilitate primer design; Clustal Omega [25], K-align [26], and MUSCLE [27,28] for multiple sequence alignments; Codeml [29], PAL2NAL [30], and Slr [31] for selection analysis; jModelTest2 [32] and ProtTest3 [33] to determine the best-fit model of evolution; and PhyML [34,35] for phylogeny reconstruction. The PhyML executable is also employed by jModelTest2 and ProtTest3. These specific tools were selected because they are some of the most popular choices and applications in many bioinformatics pipelines: for example, Primer3 is the most popular software (over 15,000 citations) for primer design [36]; Clustal Omega, K-align, and MUSCLE are very fast and accurate multiple sequence alignment tools that are commonly used to build robust DNA, RNA, or protein alignments [37]; Codeml is part of the PAML suite [29], and alongside PAL2NAL [30] and Slr [31] are commonly used to determine rates of selection [38,39]. ProtTest3 and jModelTest2 are widely used to determine best-fit models of amino-acid replacements and nucleotide substitution [40–42]; PhyML is also a popular program (over 12,000 citations) for building phylogenies using maximum likelihood [34]; for next generation sequencing data, Trimmomatic and Bowtie2 are commonly implemented in read processing and mapping pipelines [43]. Sample input files for all applications deployed by phylo-node can be found in the *Input_examples* directory and sub-directories. Taken together, phylo-node provides a diverse toolkit that allows the user to develop robust pipelines and instances using Node.

Phylo-Node is highly scalable and customizable, and was inspired by projects such as BioPerl [44] which provides very diverse tools that include Perl modules for many bioinformatic tasks and also parsers and wrappers for diverse sequence formats and applications. BioPerl’s open source structure and architecture allows users to plug new modules into BioPerl pipelines to design new applications. Node.js implements prototypal inheritance as per JavaScript but also provides access to the *module.exports* object which permits easy portability between the phylo-node toolkit and any other modules, and also interoperation between different languages by using the *child.process.exec* process. Therefore, *phylo-node* can be integrated with existing Node.js bioinformatics tools [45,46] or software written in other languages. For example, jModelTest2, ProtTest3 and Trimmomatic require a Java runtime environment (http://www.oracle.com/technetwork/java/javase/downloads/jre8-downloads-2133155.html), and by using *require* to import each module, the user can execute the analysis of these tools.

## CONCLUSIONS

In conclusion, phylo-node is a novel package that leverages the speed of Node.js to provide a robust and efficient toolkit for researchers conducting molecular phylogenetics. phylo-node can be easily employed to develop complex but consistent workflows, and integrated with existing bioinformatics tools using the Node.js codebase.

## AVAILABILITY AND REQUIREMENTS

- Project name: phylo-node
- Project home page: https://github.com/dohalloran/phylo-node
- Operating system(s): Platform independent
- Programming language: Node.js
- Other requirements: none
- License: MIT
- Any restrictions to use by non-academics: no restrictions or login requirements

## ACKNOWLEDGMENTS

I thank members of the O’Halloran lab for critical reading of the manuscript, and would like to thank The George Washington University (GWU) Columbian College of Arts and Sciences, GWU Office of the Vice-President for Research, and the GWU Department of Biological Sciences for Funding.

## AUTHOR CONTRIBUTIONS

D.O’H. conceived the idea for *phylo-node*, wrote and tested the code, and wrote the manuscript.

